# Non-permissive SARS-CoV-2 infection in human neurospheres

**DOI:** 10.1101/2020.09.11.293951

**Authors:** Carolina da S. G. Pedrosa, Livia Goto-Silva, Jairo R. Temerozo, Leticia R. Q. Souza, Gabriela Vitória, Isis M. Ornelas, Karina Karmirian, Mayara A. Mendes, Ismael C. Gomes, Carolina Q. Sacramento, Natalia Fintelman-Rodrigues, Vinicius Cardoso Soares, Suelen da Silva Gomes Dias, José Alexandre Salerno, Teresa Puig-Pijuan, Julia T. Oliveira, Luiz G. H. S. Aragão, Thayana C. Q. Torquato, Carla Veríssimo, Diogo Biagi, Estela M. Cruvinel, Rafael Dariolli, Daniel R. Furtado, Helena L. Borges, Patrícia T. Bozza, Stevens Rehen, Thiago Moreno L. Souza, Marília Zaluar P. Guimarães

**Author notes:** These authors contributed equally to this work (Carolina da S. G. Pedrosa, Livia Goto-Silva, Jairo R. Temerozo). These authors also contributed equally to this work (Stevens Rehen, Thiago Moreno L. Souza, Marília Zaluar P. Guimarães). Corresponding authors (emails) (SR), (TMLS) and (MZPG).

## Abstract

Coronavirus disease 2019 (COVID-19) was initially described as a viral infection of the respiratory tract. It is now known, however, that several other organs are affected, including the brain. Neurological manifestations such as stroke, encephalitis, and psychiatric conditions have been reported in COVID-19 patients, but the neurotropic potential of the virus is still debated. Herein, we sought to investigate SARS-CoV-2 infection in human neural cells. We demonstrated that SARS-CoV-2 infection of neural tissue is non-permissive, however, it can elicit inflammatory response and cell damage. These findings add to the hypothesis that most of the neural damage caused by SARS-CoV-2 infection is due to a systemic inflammation leading to indirect harmful effects on the central nervous system despite the absence of local viral replication.

## 1. Introduction

Severe acute respiratory syndrome coronavirus 2 (SARS-CoV-2) is the causative agent of the 2019 coronavirus disease (COVID-19), an airborne infectious disease. Most affected patients have symptoms of respiratory infections, such as fever, dry cough, and dyspnea, and 5 to 10% of them evolve to severity/death. Respiratory failure is the main event that explains fatality in COVID-19. Some authors raised the possibility that at least part of the respiratory manifestations caused by COVID-19 could be due to direct viral damage to the respiratory center located in the brain stem (Gupta et al., 2020).

Apart from respiratory symptoms, anosmia and ageusia were identified as hallmarks of COVID-19. Indeed, evidence of neurological disease associated with COVID-19 is increasing; 30 to 60% of patients further present one or more neurological symptoms, including paresthesia, altered consciousness, and headache (De Felice et al., 2020; Wang et al., 2020). Besides, COVID-19 patients are at a 7-fold higher risk to suffer a stroke, and 2-6% progress to cerebrovascular disease (Fifi and Mocco, 2020). Less frequent neurological manifestations include encephalopathy, encephalitis and Guillain-Barré syndrome (Ellul et al., 2020; Mao et al., 2020). In addition, there is mounting evidence for neurological sequelae following recovery from COVID-19, such as cognitive decline and cortical hypofunction (Hosp et al., 2021).

Neuroinvasive capacity was shown for other highly pathogenic coronaviruses, including the Middle East Respiratory Syndrome coronavirus (MERS-CoV) (Li et al., 2016) and SARS-CoV (Glass et al., 2004; Xu et al., 2005). Similarly to SARS-CoV, SARS-CoV-2 uses the angiotensin-converting enzyme 2 (ACE2) receptor as the main cellular entry (Yan et al., 2020). ACE2 is highly expressed in nasal epithelial cells, which underlies the initiation of respiratory tract infection (Sungnak et al., 2020), but is also present in multiple tissues (Hamming et al., 2004). Studies dissecting the putative mechanisms whereby SARS-CoV-2 enters the central nervous system (CNS) are still scarce. At first, it was speculated that the virus infects the olfactory neurons and reaches the brain by anterograde axonal transport. However, it was shown in rodents that SARS-CoV-2 infects non-neural cells in the olfactory epithelium rather than olfactory neurons, which could be attributed to the lack of expression of ACE2 in these cells (Brann et al., 2020; Bryche et al., 2020). In contrast, a recent study found viral particles throughout olfactory neuron projections and in the olfactory bulb of a few patients (Meinhardt et al., 2020) and other hypotheses have been proposed (Barrantes, 2021). Therefore, due to these conflicting findings, it is still debatable whether the mechanism of SARS-CoV-2 invasion in the CNS is via olfactory neurons.

Viral infections targeting the brain were successfully modeled *in vitro* using human neurospheres and brain organoids (Garcez et al., 2016; Qian et al., 2016). These models are comprised of neural cells in a tissular arrangement more like the CNS, which support more complex cellular interactions and consequently more reliable responses to viral infection. Because of that, these models became valuable for investigating SARS-CoV-2 infection in the brain tissue. While some researchers found that the virus infects neurons and glia (Jacob et al., 2020a; Mesci et al., 2020; Song et al., 2020), or a small subset of neurons (Dobrindt et al., 2021), others were unable to detect viral antigens in these cells (Pellegrini et al., 2020), suggesting SARS-CoV-2 neural infection was dependent on experimental conditions and/or viral isolates. Additionally, even fewer studies measured whether infection led to the production of new infective particles, or virions.

Here, we investigated SARS-CoV-2 infection in human neurospheres (NP) derived from induced pluripotent stem cells. Although viral infection was found to be non-permissive, it led to inflammatory response and cytotoxicity which was correlated to the amount of virus in the inoculum. We suggest that despite insufficient viral infection and replication in the brain parenchyma, there are consequences of the viral presence in the tissue, which may be enhanced by the entry of inflammatory molecules resulting from a generalized inflammatory response.

## 2. Materials and methods

### 2.1. Cell cultivation

Human induced pluripotent stem (iPS) cells were obtained from the Coriell Institute for Medical Research repository (GM23279A) or produced in house with CytoTune™-iPS 2.0 Sendai Reprogramming Kit (A16517-Invitrogen) from skin fibroblasts (Da Silveira Paulsen et al., 2012) or urine epithelial cells (Sochacki et al., 2016) (1 donor for each cell type) (Table 1). iPSC were cultured in StemFlex media (A33494 - Thermo Fisher Scientific) on top of Matrigel (BD Biosciences, Franklin Lakes, NJ). Neural Stem Cells (NSCs) were generated from iPSC following the published Life Technologies protocol for “Induction of Neural Stem Cells from Human Pluripotent Stem Cells” (Publication number: MAN0008031). NSCs were cultured in neural expansion medium (Advanced DMEM / F12 and Neurobasal medium 1:1 with 1% neural induction supplement) on a surface previously coated with Geltrex LDEV-Free (Thermo Fisher Scientific, A1413302) for expansion.

**Table 1:**
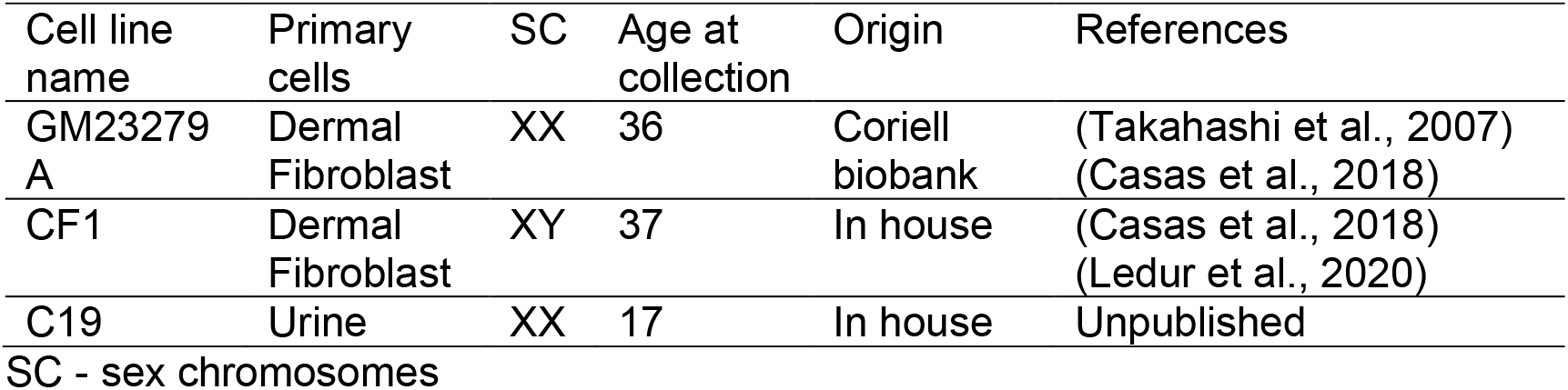
Cell lines used to generate human neurospheres.

| Cell line name | Primary cells | SC | Age at collection | Origin | References |
| --- | --- | --- | --- | --- | --- |
| GM23279 A | Dermal Fibroblast | XX | 36 | Coriell biobank | (Takahashi et al., 2007)<br>(Casas et al., 2018) |
| CF1 | Dermal Fibroblast | XY | 37 | In house | (Casas et al., 2018)<br>(Ledur et al., 2020) |
| C19 | Urine | XX | 17 | In house | Unpublished |

SC - sex chromosomes

Neurospheres (NP) were prepared in suspension as follows: NSCs were split with Accutase (Merck Millipore, SCR005) and 3 × 10^6^ cells (9 × 10^2^ cells/NP) were plated in a 6 well-plate, being cultured on an orbital shaker at 90 rpm for 7 days in standard culture conditions. At day 3, the culture medium was changed to induce differentiation, containing Neurobasal medium (Thermo Fisher Scientific, 12348017)/DMEM-F12(Life Technologies, 11330-032) 1:1, supplemented with 1x N2 (Thermo Fisher Scientific, 17502001) and 1x B27 (Thermo Fisher Scientific, 17504001), and it was changed each 3-4 days (Casas et al., 2018). Alternatively, NP were prepared directly in 96 well plates. NSCs were split with Accutase (Merck Millipore, SCR005), centrifuged at 300 x*g* for 5 minutes and resuspended in the same differentiation medium described for the 6 well-plate NP. Subsequently, 9 × 10^3^ cells in 150 μL were plated per well of a round bottom Ultra-low attachment 96-well plate (Corning, 7007), followed by centrifugation at 300 x*g* for 3 minutes to assure sedimentation. To minimize neurosphere damage, the medium of each well was changed every other day by only removing 70-100 μL and adding 150 μL of new medium. iPSC-derived human cardiomyocytes were purchased from Pluricell (São Paulo, Brazil) (Cruvinel et al., 2020). CMs were used between day 30 and day 40 of differentiation. Cell differentiation was determined by staining for cardiac specific troponin T protein and quantification by flow cytometry and found to be 86 ± 6.2%.

### 2.2. Total RNA isolation

NP were cultured in 6-well plates for 7 days, harvested and immediately frozen at -80°C until further processing. Total RNA isolation was performed using ReliaPrep^™^ RNA Tissue Miniprep System (Promega Corporation) according to manufacturer’s instructions. For cardiomyocytes, total RNA was isolated using TRIzol^™^ reagent, according to manufacturer’s recommendations (Thermo Fisher Scientific). RNA concentration and quality were quantified on a NanoDrop^™^ 2000c Spectrophotometer (Thermo Fisher Scientific); and integrity and purity were evaluated by 1.8% agarose gel electrophoresis using a photo documentation device equipped with a UV lamp (L-PIX, Loccus Biotecnologia). Then, samples were digested with DNase I, Amplification Grade, following the manufacturer’s instructions (Invitrogen, Thermo Fisher Scientific). 2 μg of RNA from DNAse-treated samples were reverse transcribed using M-MLV for complementary DNA generation (cDNA) (Thermo Fisher Scientific).

### 2.3. Quantitative Reverse Transcriptase–Polymerase Chain Reaction (qRT-PCR)

For gene expression analysis of NP and cardiomyocytes, qRT-PCR reactions were conducted in three replicates with a final reaction volume of 10 µL in MicroAmp Fast Optical 96 Well Reaction Plates (Thermo Fisher Scientific) containing 1X GoTaq qPCR Master Mix (Promega Corporation), 300 nM CXR Reference Dye, a final concentration of 200 nM of each SYBR green designed primers [Angiotensin I Converting Enzyme 2 (ACE2; forward: 5’-CGAAGCCGAAGACCTGTTCTA-3’; reverse: 5’-GGGCAAGTGTGGACTGTTCC-3’) Thermo Fisher Scientific] and 10 ng of cDNA per reaction. Appropriate controls (no reverse transcriptase and template-negative controls) were incorporated into each run. Briefly, the reactions were performed on a StepOnePlus™ Real-Time PCR System thermocycler (Applied Biosystems). Thermal cycling program comprised of a denaturing step at 95°C for 3 min, followed by 40 cycling stages at 95°C for 15 sec, 57°C for 15 sec, 72°C for 15 sec and melt curve stage 95 °C, 15 sec; 60 °C, 1 min; 95 °C, 15 sec. The relative expression of the genes of interest was normalized by human reference genes: Glyceraldehyde-3-phosphate Dehydrogenase (GAPDH; forward: 5’-GCCCTCAACGACCACTTTG-3’; reverse: 5’-CCACCACCCTGTTGCTGTAG-3’) and Hypoxanthine Phosphoribosyltransferase 1 (HPRT-1; forward 5’-CGTCGTGATTAGTGATGATGAACC-3’; reverse: 5’-AGAGGGCTACAATGTGATGGC-3’). qPCR data analysis was performed with the N_0_ method implemented in LinRegPCR v. 2020.0, which considers qPCR mean efficiencies estimated by the window-of-linearity method (Ramakers et al., 2003; Ruijter et al., 2009). Briefly, N_0_ values were calculated in LinRegPCR using default parameters. Then, the arithmetic mean of N_0_ values from gene of interest (GOI) was normalized by taking its ratio to the N_0_ geometric mean of the reference genes (REF: *GAPDH and HRRT-1*; N_0GOI_/N_0REF_).

### 2.4. SARS-CoV-2 propagation

SARS-CoV-2 obtained from a nasopharyngeal swab from a confirmed case in Rio de Janeiro, Brazil (GenBank accession no. MT710714, PANGO lineage B.1 (Rambaut et al., 2020), clade 20C (Hadfield et al., 2018) was expanded in African green monkey kidney cells (Vero, subtype E6). Virus isolation was performed after a single passage in cell culture in a 150 cm^2^ flask, previously infected with SARS-CoV-2 at multiplicity of infection (MOI) 0.01. All procedures related to virus culture were handled in a biosafety level 3 (BSL3) facility, according to WHO guidelines. Virus stocks were kept at -80°C.

### 2.5. Cell infection

NP cultivated for 7 days were infected with SARS-CoV-2 at MOI 0.1 and 0.01. Following 1h incubation, the inoculum was partially replaced by fresh medium and NP were cultivated for additional 2 and 5 days, with orbital shaking at 90 rpm in standard conditions (5% CO_2_ and 37°C). The medium was not completely replaced to avoid excessive stress as explained above. NP exposed to uninfected Vero’s culture medium were used as controls of infection (Mock). NP (≈ 50 to 200) were used per experimental group for each analysis. Culture supernatants were collected at 2- and 5-days post-infection for virus titration by plaque forming units (PFU) assay and/or real time qRT-PCR. The assay was performed in a single experiment, consisting of three cell lines generated from 3 independent donors.

In lieu of the formerly described method, seven days-old NP were separately (one per well) infected for 1h or 24h with SARS-CoV-2 at different MOIs (0.1, 1 and 10). After incubation, the inoculum was partially replaced by fresh medium, for the reasons stated above, and NP were cultured in standard conditions (5% CO_2_ and 37°C). Supernatants and NP were collected at 24h, 48h and 72h post-infection. NP exposed to uninfected Vero’s culture medium were used as controls of infection (Mock). 5 to 24 NP were distributed by each experimental group.

Cardiomyocytes were infected with SARS-CoV-2 at MOI 0.1 for 1h. Next, the inoculum was replaced by fresh medium and cultured in standard conditions for 48-72 h. After that, the monolayer was fixed with 4% paraformaldehyde (PFA) solution.

### 2.6. Immunofluorescence staining

NP were fixed in 4% paraformaldehyde solution (Sigma-Aldrich) for 1h, followed by cryopreservation with 30% sucrose solution overnight. Then, samples were embedded in O.C.T compound (Sakura Finetek, Netherland) and frozen at -80°C. The O.C.T blocks were sectioned in 20 μm slices with a Leica CM1860 cryostat. After washing with PBS, sections were incubated in permeabilization/blocking solution (0.3% Triton X-100/ 3% goat serum) for 2h. The primary antibody was incubated overnight at 4°C [anti-double-stranded RNA (dsRNA) monoclonal antibody (1:200, Scicons)]. Then, sections were incubated with secondary antibody goat anti-mouse Alexa Fluor 488; 1:400, A-11001 - Thermo Fisher Scientific) for 2h. Nuclei were counterstained with 0.5 μg/mL 40-6-diamino-2-phenylindole (DAPI) for 10 minutes and the slides were mounted with Aqua-Poly/Mount (Polysciences).

After fixation, cardiomyocytes were washed with PBS and then incubated with permeabilization/blocking solution (0.3% Triton X-100 / 3% bovine serum albumin) for 1h. Then, cells were incubated with primary antibody anti-dsRNA (1:200) overnight at 4°C. On the next day, cells were incubated with the secondary antibody goat anti-mouse Alexa Fluor 488 (1:400) for 1h. Nuclei were counterstained with DAPI for 5 minutes and mounted with 50% PBS-Glycerol.

Images of NP and cardiomyocytes were acquired on a Leica TCS-SP8 confocal microscope with the 63x and 20x objective, respectively.

### 2.7. Flow Cytometry

CM were plated on Geltrex™ (Thermo Scientific)-coated 6-well plates up to days 30-32 of differentiation. Then, cells were dissociated, fixed with 1% paraformaldehyde solution and permeabilized with 0.1% Triton (Sigma Aldrich) and 0.1% Saponin (Sigma Aldrich). CM were stained with anti-TNNT2 (1:2500, MA5-12960 - Thermo Scientific). The data of each batch of differentiation was acquired using a Canto BD Flow cytometer and analyzed using the FlowJo Software considering 1%–2% of false-positive events. iPSC were used as negative control for the reaction for definitive CM. We used samples incubated only with secondary antibodies to draw the limits of positive events.

### 2.8. Identification and quantification of neural progenitor population

NP were fixed, cryopreserved, embedded in O.C.T compound and frozen as described in the immunofluorescence staining section. After cryosectioning at 20 μm, NP were washed in PBS and incubated in permeabilization solution (0.3% Triton X-100 in PBS) for 20 min followed by incubation in blocking solution (3% bovine serum albumin/ 5% goat serum) for 2h. The primary antibody was incubated overnight at 4°C [human Pax6 polyclonal antibody (1:100, Thermo Fisher Scientific)]. Then, sections were incubated in blocking solution, as described previously; followed by secondary antibody goat anti-rabbit Alexa Fluor 488; 1:400, A-11008 - Thermo Fisher Scientific) for 2h. Nuclei were counterstained with 0.5 μg/mL 40-6-diamino-2-phenylindole (DAPI) for 10 minutes and slides were mounted with Aqua-Poly/Mount (Polysciences).

Images of NP were acquired on a Leica TCS-SP8 confocal microscope with the 63x objective and analyzed with Image J software. PAX6+ area was normalized over nuclei staining area. 10 NP per experimental group were analyzed for each cell line, totalling 20 NP from 2 different cell lines (CF1 and C19). Statistical analysis to compare differences among groups was performed using One way ANOVA with Tukey’s post-hoc test (significance set at p<0.05).

### 2.9. Plaque forming unit assay

For virus titration, monolayers of Vero E6 cells (2 × 10^4^ cell/well) in 96-well plates were infected with serial dilutions of supernatants containing SARS-CoV-2 for 1 hour at 37°C. Semi-solid high glucose DMEM medium containing 2% FBS and 2.4% carboxymethylcellulose was added and cultures were incubated for 3 days at 37 °C. Then, the cells were fixed with 10% formalin for 2 h at room temperature. The cell monolayer was stained with 0.04% solution of crystal violet in 20% ethanol for 1 h. Plaque numbers were scored in at least 3 replicates per dilution by independent readers blinded to the experimental group and the virus titers were determined by plaque-forming units (PFU) per milliliter.

### 2.10. Molecular detection of SARS-CoV-2 RNA

The total RNA from NP’s supernatant was extracted using QIAamp Viral RNA (Qiagen), according to manufacturer’s instructions. Quantitative RT-PCR was performed using QuantiTect Probe RT-PCR Kit (Quiagen®) in a StepOne™ Real-Time PCR System (Thermo Fisher Scientific). Amplifications were carried out in 25 µL reaction mixtures containing 2X reaction mix buffer, 50 µM of each primer, 10 µM of probe, and 5 µL of RNA template. Primers, probes, and cycling conditions recommended by the Centers for Disease Control and Prevention (CDC) protocol were used to detect the SARS-CoV-2. Alternatively, genomic (ORF1) (gRNA) and subgenomic (ORF(sgRNAE)) were detected, as previously described (Wölfel et al., 2020). In summary, for detection of gRNA, the following primers and probe targeting for sequences downstream of the start codons of the gene E were used (E_Sarbeco_F; ACAGGTACGTTAATAGTTAATAGCGT / E_Sarbeco_R; ATATTGCAGCAGTACGCACACA / E_Sarbeco_P1; FAM-ACACTAGCCATCCTTACTGCGCTTCG-BBQ); and for sgRNA detection, a leader-specific forward primer (sgLeadSARSCoV2-F; CGATCTCTTGTAGATCTGTTCTC) was used with the reverse primer and probe for gene E above mentioned, and described in (Corman et al., 2020; Wölfel et al., 2020).

### 2.11. Western blot analysis

Two days post-infection (d.p.i), 100 µL of sample buffer without bromophenol blue (62.5 mM Tris-HCl, pH 6.8, containing 10% glycerol, 2% SDS and 5% 2-mercaptoethanol) was added to the NP and then, samples were frozen at -80°C. Next, samples were gently broken down with a disposable pestle (BAF 199230001-Sigma, Bel-Art) and cell extracts were boiled at 95°C for 10 min and centrifuged at 4°C 16,000x g for 15 min to remove insoluble material. Protein content was estimated using the Bio-Rad Protein Assay (#5000006, Biorad). After addition of bromophenol blue (0.02%), extract samples (40 µg/lane for NP and 15 µg/lane for Vero cells) were separated by electrophoresis on a 10% SDS polyacrylamide gel and transferred to polyvinylidene difluoride (PVDF) membranes. Membranes were blocked in 5% non-fat milk in Tris-Buffered Saline with 0.1% Tween-20 (TBS-T) for 1 hour at room temperature. Membranes were then incubated overnight at 4°C, in the presence of anti-SARS-CoV-2 SP (1:2,000, #GTX632604 - GeneTex) and anti-actin (1:2000, MAB1501, Millipore) diluted in TBS-T with 5% non-fat milk. Then, membranes were incubated with peroxidase-conjugated antibody goat anti-Mouse IgG (H+L), HRP-conjugate (1:10,000, G21040 -Molecular Probes). The signals were developed using ECL Prime Western Blotting System (#GERPN2232, Sigma) for five minutes and chemiluminescence was detected with an Odyssey-FC System® (Imaging System - LI-COR Biosciences). Membranes were also stripped for re-staining by incubating for three cycles of 10 minutes in stripping buffer (pH 2.2, 200 mM glycine, SDS 0,1% and 1% Tween-20), the buffer was discarded, then the membranes were washed for 5 minutes with PBS (three times) and 5 minutes with 0.1% TBS-T (three times). Next, the membranes were blocked again and processed for immunolabeling as described above.

### 2.12. Cytokine multiplex assay and LDH cytotoxicity assay

A multiplex biometric immunoassay containing fluorescent dyed microbeads was used to measure cytokines in the cell culture supernatant (Bio-Rad Laboratories, Hercules, CA, USA). The following cytokines were quantified: IL-1β, IL-1RA, IL-2, IL-4, IL-5, IL-6, IL-7, IL-8, IL-9, IL-10, IL-12(p70), IL-13, IL-15, IL-17, basic FGF, Eotaxin, G-CSF, GM-CSF, IP-10, MCP-1, MIP-1α, MIP-1β, PDGF-BB, RANTES, TNF-α, and VEGF; and cytokine levels were calculated by Luminex technology (Bio-Plex Workstation; Bio-Rad Laboratories, USA). The analysis of data was performed using software provided by the manufacturer (Bio-Rad Laboratories, USA). A range of 0.51–8,000 pg/mL recombinant cytokines was used to establish standard curves and the sensitivity of the assay. Cytotoxicity was determined according to the activity of lactate dehydrogenase (LDH) in the culture supernatants using a CytoTox® Kit (Promega, USA) according to the manufacturer’s instructions, and presented as OD (optical density) of infected cells minus OD of mock. Statistics were performed using GraphPad Prism software version 8. Numerical variables from NP experiments were tested regarding their distribution using the Shapiro-Wilk test. For those following a normal (parametric) distribution, One-way analysis of variance (ANOVA) with Dunnet’s post-hoc test was used to compare differences among groups; and for nonparametric data, Kruskal-Wallis test with Dunn’s post-hoc test was used to compare differences.

### 2.13. SARS-CoV-2 strains sequence analyses

Nucleotide sequences from our strain (GenBank accession No. MT710714), from strains used in others works (Song et al., 2020; Zhang et al., 2020), and from SARS-CoV-2 reference genome (Wuhan-Hu-1, GenBank accession No. NC_045512.2) were retrieved from NCBI database and from Multiple Sequence Alignment (MSA). Visualization of SARS-CoV-2 sequences was performed using ClustalW (Thompson et al., 1994) implemented in MEGA X program-version 10.1.8 (Kumar et al., 2018). Maximum Likelihood phylogenetic analyses were conducted using the JTT matrix-based model (Jones et al., 1992) with confidence assessed by bootstrap with 1,000 replicates. For identification of amino acid mutations, the coding data was translated assuming a Standard genetic code and compared to the SARS-CoV-2 reference genome Wuhan-Hu-1.

## 3. Results

### 3.1. SARS-CoV-2 replication in human neural cells is non-permissive

Histopathological analyses of several tissues from an infant deceased with COVID-19 fully described in a case report from our group (Gomes et al., 2020), showed viral presence by immunofluorescence (IF) with anti-SARS-CoV-2 spike protein (SP) in the choroid plexus (ChP), lateral ventricle (LV) lining, and in some locations of the frontal cerebral cortex (S1 Fig). SARS-CoV-2 infection was confirmed by quantitative Reverse Transcriptase–Polymerase Chain Reaction (qRT-PCR) (S1 Table). The heart and the lungs, which have been previously demonstrated to be permissive to SARS-CoV-2 infection (Gnecchi et al., 2020; Pérez-Bermejo et al., 2020; Schaller et al., 2020; Tabary et al., 2020), also showed presence of the virus (S1 Table and (Gomes et al., 2020)). We sought to verify whether the expression levels of the ACE2 receptor, which could be retrieved from Protein Atlas and Allen Brain Atlas databases (Hawrylycz et al., 2012; Uhlén et al., 2015) could correlate to the differences in viral infection in different brain areas. These databases contain RNAseq levels from donors between 0 and 96 years-old (Protein Atlas) and 24 and 57 years-old (Allen Brain Atlas). For comparison, we also looked at the expression levels in the heart and lungs, which are known to be infected by SARS-CoV-2. We found that ChP had higher ACE2 levels than other brain regions assessed (Fig 1) and the heart and lungs expressed higher levels of ACE2 in comparison with most neural tissues (Fig 1A).

**Fig 1.**
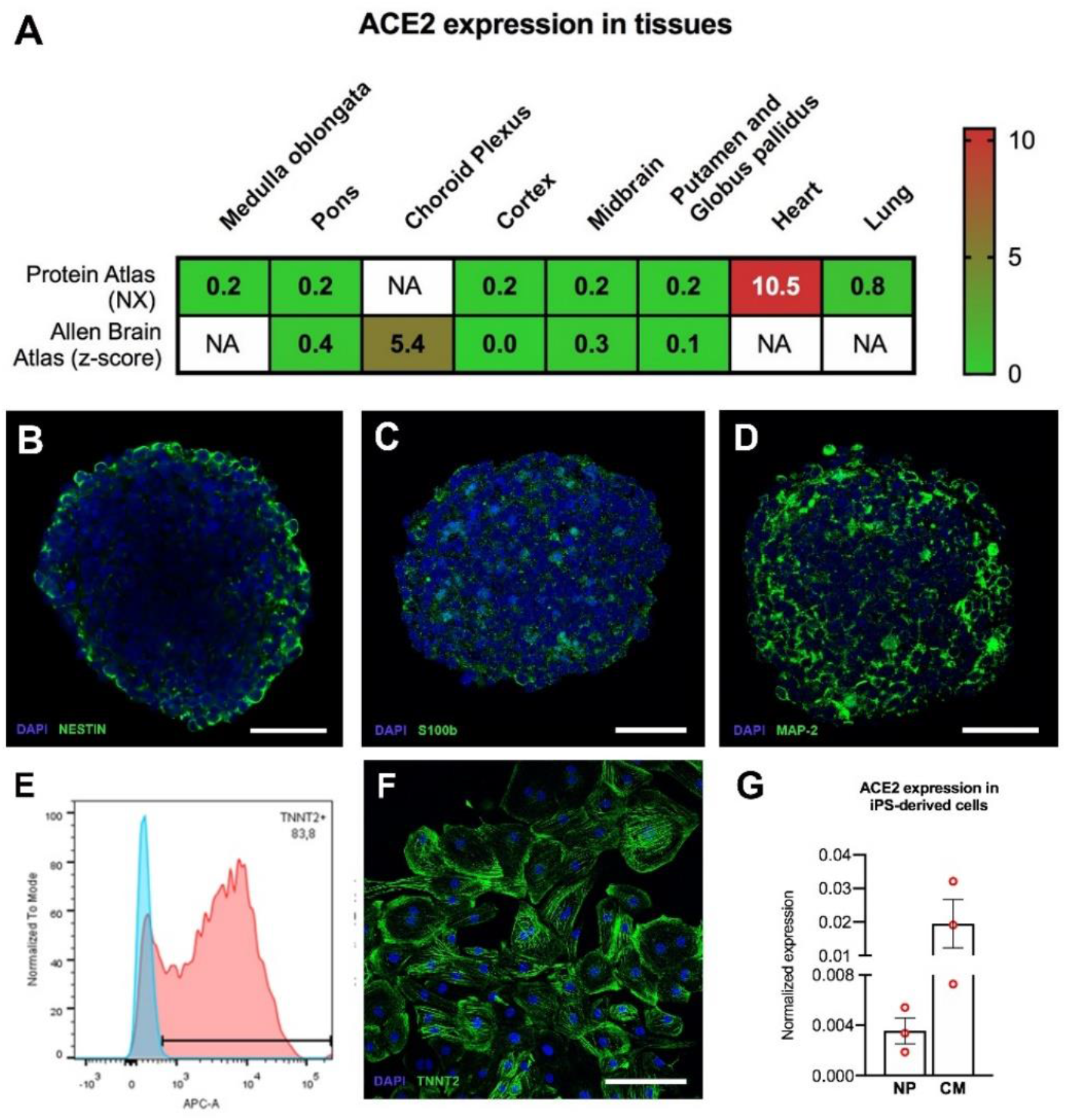
ACE2 expression in adult human tissues and iPSC-derived cells. (A) ACE2 expression in adult human tissues based on Protein Atlas and Allen Brain Atlas databases. Consensus normalized expression (NX) and z-score are used to represent the relative abundances within the respective databases (Citations in the main text). NA= not analyzed in the database. Characterization of NP and CM. NP were cultivated for 7 days and CM were differentiated for 35 days and characterized by IF (B-D and F) and flow cytometry (E). (B) NP were immunostained for Nestin (neural progenitors), (C) S100b, (astrocytes); and (D) MAP-2 (neurons). (E) Representative flow cytometry graph showing a quantification of 83.2% TNNT2+ cells from one of the cell batches used in this study; (F) CM immunostained for cardiac troponin T (TNNT2). Scale bars: (B-D) 40 µm and (F) 250 µm. (G) Relative mRNA expression levels of ACE2 in iPSC-derived cultures; NP cultivated for 7 days and CM differentiated for 35 days. (n=3 cell culture batches for both), normalized to reference genes, GAPDH and HPRT-1. The data correspond to 3 independent cell lines for NP.

NP have been previously used to investigate ZIKV infection and in studies to address drug neurotoxicity and neural activity (Garcez et al., 2017, 2016; Pampaloni et al., 2007; Sirenko et al., 2019; Woodruff et al., 2020). NP are obtained from neural stem cells (NSCs) cultured in suspension to form cellular aggregates in the presence of neural differentiation medium. This process triggers the differentiation of NSCs into neural cell types including neurons and astrocytes. Neuronal, astrocytic and neural progenitor proteins have been identified in a proteomic analysis of neurospheres recently published by our group (Goto-Silva et al., 2021). Here, a characterization of NP shows immunostaining for neurons (MAP-2), astrocytes (s100b) and neural progenitors (Nestin) (Fig 1B-D). Human iPSC-derived CM express the ACE2 receptor and were used as an *in vitro* model for cardiac SARS-CoV-2 infection (Marchiano et al., 2020; Pérez-Bermejo et al., 2020). The CMs presently used were mostly TNNT2 positive (86 ± 6.2% positive cells at 30-32 days of differentiation) (Fig 1E and 1F).

Then we checked whether human NP and cardiomyocytes (CM) would mirror ACE2 levels of brain and, consequently, serve the purpose of modeling SARS-CoV-2 infection. ACE2 expression in NP was, on average, almost one order of magnitude lower than in CMs. (Fig 1G). This shows that indeed NP reproduce the differences in tissue expression of ACE2 and validates the use of NP as a neural model of SARS-CoV-2 infection *in vitro*. Interestingly, NSCs express much less ACE2 and CD147, whereas AXL is unchanged, when compared to NP (S2B Fig), the former two well-established SARS-CoV-2 receptors. These differences suggest that neuroprogenitors may be less susceptible to SARS-CoV-2 infection than differentiated neural cells.

Having established the viability of these models, we then exposed NP and CM to the new coronavirus. SARS-CoV-2-infection in NP was addressed using multiplicity of infection (MOI) 0.01 and 0.1 and a 1h exposure to viral particles, all conditions known to infect other virus-permissive cells, such as CM, Vero and Calu-3 cells (Han et al., 2020; Lamers et al., 2020; Marchiano et al., 2020; Pérez-Bermejo et al., 2020). Like other previous works, at first we evaluated SARS-CoV-2 infection using immunodetection of viral macromolecules (Dobrindt et al., 2021; Jacob et al., 2020b; Song et al., 2021; Zhang et al., 2020). At five days post infection (d.p.i.), SARS-CoV-2 could not be detected by immunostaining for double-stranded (ds)RNA (Fig 2A). Additionally, infected NP did not stain with anti-SARS-CoV-2 convalescent sera (CS) (S3A Fig). In contrast, NP infected with Zika virus (ZIKV) were immunoreactive for dsRNA (Fig 2A and inset), and CM infected with SARS-CoV-2 were immunoreactive for dsRNA and CS (arrows, Fig 2A and S3B Fig, correspondingly). The absence of immunostaining for SARS-CoV-2 was surprising because 5 dpi it was expected that the virus would have replicated. At an earlier time point, 2 dpi, we also failed to detect viral proteins, using both anti-SARS-CoV-2 SP (Fig 2B) and anti-SARS-CoV-2 CS (S3C Fig) with Western blotting (WB). On the other hand, infected Vero cells showed prominent bands corresponding to the molecular weights of SARS-CoV-2 spike protein and nucleoprotein, as detected by anti-SARS-CoV-2 SP (Fig 2B) and anti-SARS-CoV-2 CS (S3C Fig), respectively.

**Fig 2.**
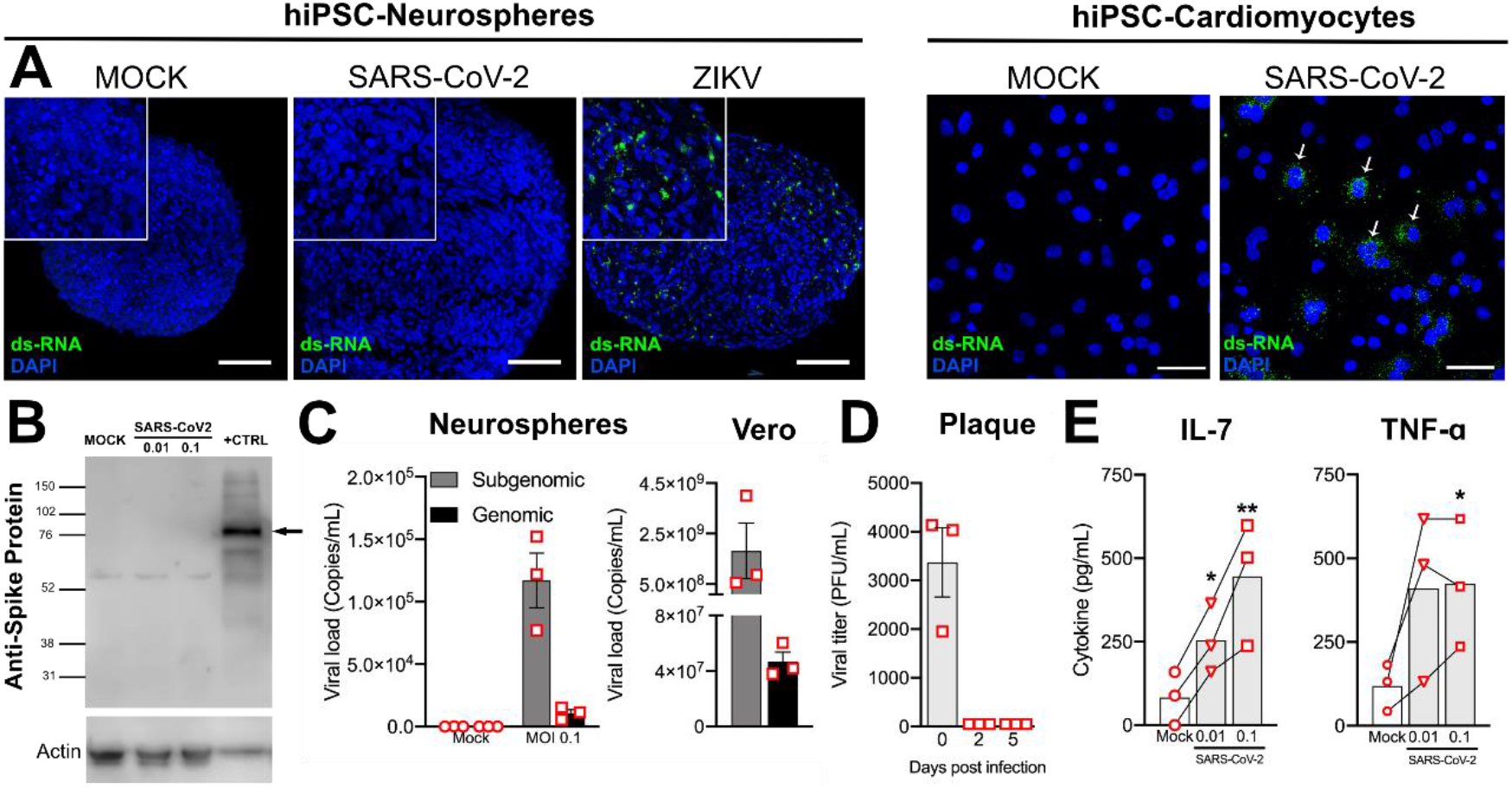
SARS-CoV-2 replication in NP is non-productive. (A) IF staining for SARS-CoV-2, with anti-dsRNA (green), in cryosections of 5 d.p.i. NP (MOI 0.1). Nuclei were counterstained with DAPI (blue). ZIKV-infected NP (MOI 0.5 for 2h, 3 d.p.i) and SARS-CoV-2-infected CM (MOI 0.1 for 1h, 2 d.p.i) were used as positive controls for dsRNA labeling and SARS-CoV-2 infectivity, respectively. Scale bar: 50 µm (B) Detection of SARS-CoV-2 SP by WB. Protein extracts from Vero cells (MOI 0.1 for 24h) were used as positive control. Gel loading was assessed by beta-actin staining. (C) Real-time qRT-PCR of genomic and subgenomic RNA levels of SARS-CoV-2 in the supernatants of NP 5 d.p.i. SARS-CoV-2-infected Vero cells were used for comparison. (D) Plaque forming units’ assay from the supernatants of the NP at 2 and 5 d.p.i (MOI 0.1). Data was plotted as average plus standard error. (E) Multiplex luminex assay for IL-7 and TNF-α from the supernatant of NP collected 5 d.p.i. (\**p*<0.05, \*\**p*<0.005). Collected data corresponds to an exposure of NP from 3 different cell lines (Table 1) to SARS-CoV-2 (PANGO lineage B.1, Nextstrain clade 20C) in one experimental infection. Each donor data point corresponds to the average of a duplicate technical measurement and is connected within the different experimental groups (cell lines).

Despite the absence of SARS-CoV-2 detection, the supernatant of NP harvested 5 d.p.i. showed approximately 10^4^ and 10^5^ copies of genomic and subgenomic viral RNA, respectively, which were at least 4 orders of magnitude lower than the observed in infected Vero cells (Fig 2C). The detection of sgRNA indicates that there had been cell entry and initiation of negative-strand synthesis of the viral genome (Wölfel et al., 2020). The plaque forming unit (PFU) assay showed no infective viral progenies at 2 or 5 d.p.i in the supernatant of NP (MOI 0.1) (Fig 2D), whereas for CM at 2 d.p.i (MOI 0.1) there was on average 2.8 × 10^6^ PFU/mL (data not shown). The analysis of the virus inoculum in NP at day 0 contained on average 3 × 10^3^ PFU/mL, confirming that they were exposed to infectious virus particles (Fig 2D). The aforementioned detection of viral genomic RNA contrasts with the absence of infectious virus progeny and may indicate that neural cells were a dead-end for SARS-CoV-2 infection, or that the viral RNA detected in the supernatant was a residual of the inoculum.

### 3.2. SARS-CoV-2 infection of neural cells triggers an increase in pro inflammatory cytokines

SARS-CoV-2 infection has been associated with cytokine storm as a poor prognosis of disease progression (Tay et al., 2020). Misbalance of cytokine levels in the CNS microenvironment is associated with several pathologies (Guzman-Martinez et al., 2019; Lima et al., 2019; Mutso et al., 2020; Sochocka et al., 2017), hence the need to uncover the brain inflammation associated with SARS-CoV-2 infection. To investigate if SARS-CoV-2 generates a direct inflammatory response in neural tissues, the supernatant of infected NP was collected at 5 d.p.i. and analyzed by Multiplex Luminex assay to measure over 20 cytokines and chemokines (Fig 2E and S4 Fig). The levels of IL-7 and TNF-α in the infected NP were higher compared with mock condition (Fig 2E). The levels of other cytokines and chemokines were not affected by SARS-CoV-2 infection in these conditions (S4 Fig).

### 3.3. Prolonged exposure to higher inocula of SARS-CoV-2 does not elicit permissive infection of NP

The exposure time of NP to the virus inoculum and the low MOIs are sufficient to detect the virus in permissive cells, including Vero and CM. The same conditions, in contrast, did not prompt the detection of viral particles by neither IF nor WB in NP nor even infective particles in the culture supernatant. Arguably, this evidence of non-permissiveness could be explained by insufficient exposure time of the cells during viral inoculation. A previous report showed infection of NP after 24 h inoculation time at higher MOIs (Zhang et al., 2020). Additionally, it is speculated that the initial amount of virus in the inoculum is crucial in determining the severity of COVID-19 symptoms (Imai et al., 2020). To clarify these questions, NP were then exposed to a longer infection time and to higher MOIs. We emulated the protocols for neurospheres generation and infection with SARS-CoV-2 described by others in order to better compare our results with previously published data (Zhang et al., 2020).

We compared the capacity of SARS-CoV-2 to generate infectious viral particles in NP using 1h and 24h incubation times and MOIs of 0.1, 1 and 10. Despite longer exposure and higher viral titers, these conditions also did not support the production of novel infective particles, as measured by plaque forming units assay of the respective supernatants (Fig 3A). However, SARS-CoV-2 SP was detectable by IF in NP incubated for 24h with the virus and analyzed 24h post infection (Fig 3B). On the other hand, after 48h and 72h of the same 24h-infection, the NP had no detectable SP immunostaining, suggesting that viral proteins are only present transiently and then cleared (Fig 3B and S3D Fig). Remarkably, at a time point in which no virus could be detected in NP, LDH measurements of culture supernatants revealed increased cytotoxicity at MOI 1 and 10, irrespective of incubation time with the virus inoculum (Fig 3C). These data indicate that exposure of NP to SARS-CoV-2 at high MOIs is neurotoxic (Fig 3). Also, the incubation time with the inoculum does not seem to affect the outcome of infection, since prolonged exposure time did not increase the production of viral particles or augment the extent of cell damage.

**Fig 3.**
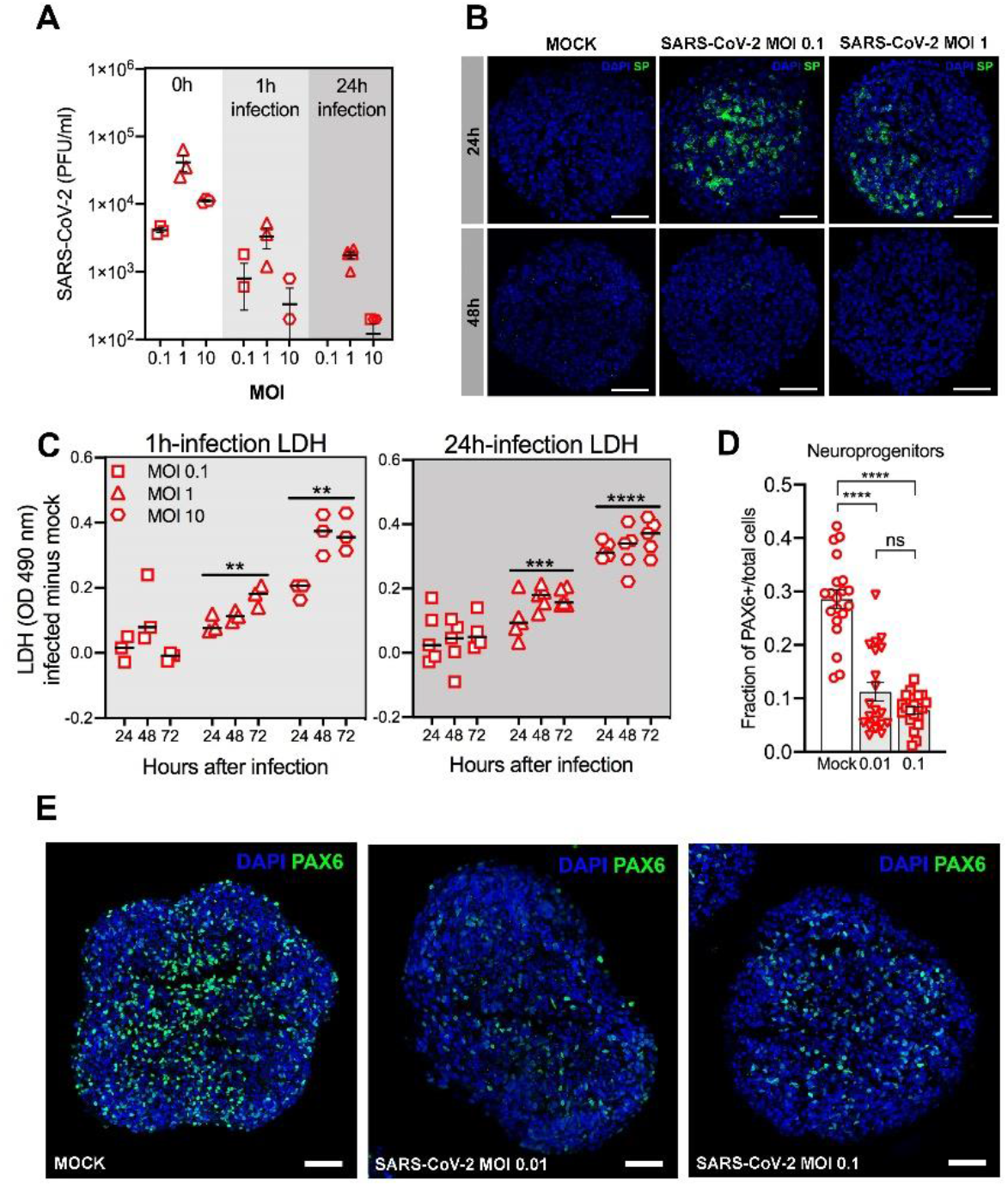
Prolonged exposure to higher inocula of SARS-CoV-2 does not elicit permissive infection of NP but higher MOIs are cytotoxic. NP were inoculated with SARS-CoV-2 for 1 h or 24 h at indicated MOIs. (A) At 72h post-infection, culture supernatants were harvested to quantify infectious virus titers in Vero cells by plaque forming units’ assay. (B) NP inoculated for 24 h were immunostained with anti-SARS-CoV-2 SP (green) and counterstained with DAPI (blue) at 24h and 48h post-infection. Scale bars: 40 µm. (C) Lactate Dehydrogenase (LDH) levels were measured by colorimetric assay in NP exposed to viral particles for 1h or 24h and analyzed at 24h, 48h and 72h post infection. Collected data correspond to the infection of 5 NP from 1 cell line, produced in house from urine epithelial cells, in one experimental infection. Each data point corresponds to measurements from single NP from one experimental infection. (D) Analysis of neural progenitor (PAX6^+^) population 5 d.p.i. at M.O.I. 0.01 and 0.1 from 2 cell lines (CF1 and C19), one experimental infection. Cryosections of NP were immunostained for PAX6 and the fraction of PAX6+ cells was quantified. Each data point represents 1 NP, and a total of 20 NP from 2 different cell lines were analyzed per group. **** p<0.0001. (E) Representative IF staining for neural progenitors (PAX6+, green), in cryosections of NP (MOI 0.01 and 0.1, 5 d.p.i.). Nuclei were counterstained with DAPI (blue). Scale bar: 50 µm.

Neural stem cells in the brain have an important role in multi-potency and self-renewal. In a study done on ocular donor tissues and iPSC-derived eye organoid model, it was observed that the limbus, an eye region containing stem cells, was particularly susceptible to infection, suggesting stemness was to blame (Makovoz et al., 2020). In order to investigate whether SARS-CoV-2 infection could affect the progenitor pool, we isolated NP nuclei by isotropic fractionation, a method previously reported to be effective to quantify cell populations in the brain (Herculano-Houzel and Lent, 2005). Using this method, the neural progenitor population in NP was identified as PAX6+ cells. We observed that SARS-CoV-2 infection did not change the proportion of neural progenitors in NP 5 days after infection but presented a trend towards progenitor pool reduction (S2A Fig). Since isotropic fractionation was done with a pool of NP which might have different degrees of infection individually, we decided to quantify the density of progenitors in single NP by immunostaining PAX6+ cells in cryosections. After quantifying about 20 NP per group from 2 different cell lines, we observed that PAX6 was significantly reduced at MOI 0.01 and 0.1 (p<0.0001), with no significant difference among the different MOIs (Fig 3D and 3E). These data suggest that infection impacts the amounts of progenitors by interfering with cell proliferation or inducing neural progenitor’s cell death. Considering that the expression of ACE2 is much lower in progenitors than in NP (S2B Fig), the mechanisms regulating reduction in progenitors during SARS-CoV-2 infection should be further explored.

### 3.4. Strain differences in SARS-CoV-2 RNA sequence could explain distinct infectivity of neural cells

Genetic variation of SARS-CoV-2 isolates has been described and could help to explain differences in the experimental infectivity of neural tissues previously reported (Korber et al., 2020). To investigate whether possible genetic components could account for the differences among SARS-CoV-2 infections reported in literature and in the current work, we compared the phylogeny of the nucleotide sequences of our strain with the strains used in these works, by means of the Wuhan strain genome as reference. The analysis showed less than 1% variation among the genetic sequences of our strain, Zhang and collaborators, and Song and collaborators (Table 2) (Song et al., 2020; Zhang et al., 2020). These small variations correlate with mutations in the receptor-binding domain of Spike protein and two non-structural proteins (nsp) (Table 3).

**Table 2.** Estimates of evolutionary divergence between SARS-CoV-2 sequences.

| **Virus Strain** | **NC_045512.2**<br>**Wuhan-01** | **MT710714.1**<br>**This study** | **MT230904.1**<br>**Zhang et al** | **MT020880.1**<br>**Song et al** | **MT246667.1**<br>**Song et al** |
| --- | --- | --- | --- | --- | --- |
| **MT710714.1 -**<br>**This study** | 0.06 | - | - | - | - |
| **MT230904.1 -**<br>**Zhang et al** | 0.03 | 0.09 | - | - | - |
| **MT020880.1 -**<br>**Song et al** | 0.01 | 0.07 | 0.04 | - | - |
| **MT246667.1 -**<br>**Song et al** | 0.01 | 0.07 | 0.04 | 0.00 |  |
| **MN985325.1 -**<br>**Song et al** | 0.01 | 0.07 | 0.04 | 0.00 | 0.00 |

The values correspond to the divergence in sequences of SARS-CoV-2 when comparing the strain in rows with the strain in columns. Sequences were compared with maximum likelihood phylogenetic analysis as described in the Material and Methods section.

**Table 3.**
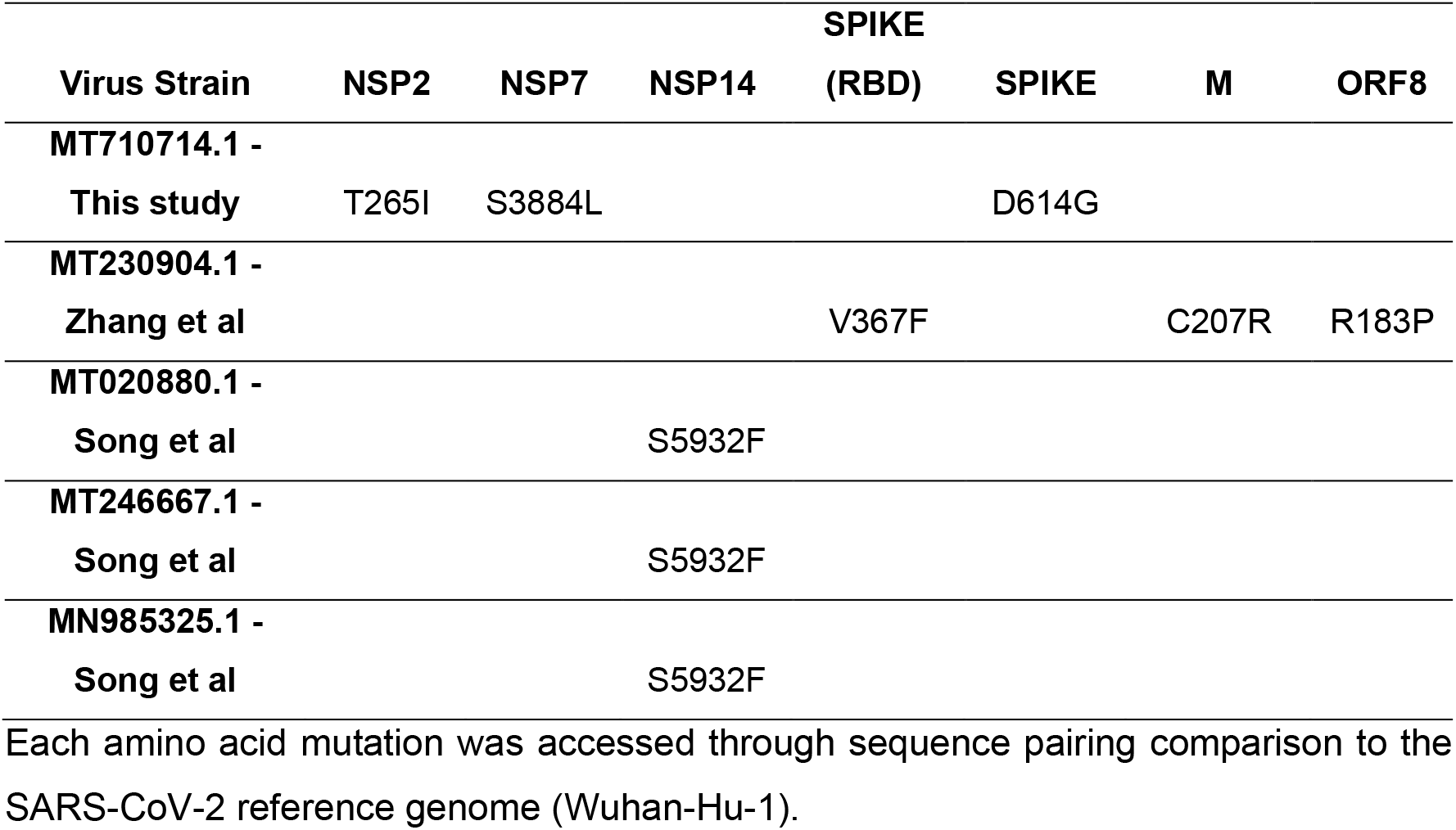
Specific proteins mutations among SARS-CoV-2 strains.

Our viral isolate presented three missense mutations compared to the Wuhan sequence: D614G on spike protein, S3884L on nsp7 and T265I on nsp2. The D614G spike protein mutation has been linked to more infectious behavior of the strain (Korber et al., 2020, p. 61). Nsp 7 participates in SARS-CoV-2 replication, being a co-factor for RNA polymerase. The nsp2 role is unknown and may be related to the disruption of intracellular host signaling and mitochondrial biogenesis (Cornillez-Ty et al., 2009).

## 4. Discussion

Neurological manifestations of COVID-19 fueled research on the putative viral infection of the brain. Absence or low detection of SARS-CoV-2 was described in the cerebrospinal fluid (CSF), and so far many reports have described the presence of SARS-CoV-2 in the brain parenchyma (Crunfli et al., 2020, p.; Espíndola et al., 2020; Schaller et al., 2020; Song et al., 2020), while others suggested quick viral clearance in the brain due to swift innate immune responses (Schurink et al., 2020).

Here, we bring the attention to SARS-CoV-2 infection in the ChP in its LV lining, and in the frontal cortex of an infant’s brain (S1A Fig), which was confirmed by qRT-PCR for viral RNA (S1 Table) (pathological analysis of this case was described in detail in (Gomes et al., 2020)).The detection of SARS-CoV-2 was more pronounced in the ChP than in the cortex (S1A Fig). This matches expression data for the receptor ACE2, which is enriched in ChP both *in vivo* and *in vitro*, in comparison to other brain areas, including cortex (Jacob et al., 2020a; Pellegrini et al., 2020) (Fig 1A). Our protocol for differentiating NP generates cortical brain-like tissue (Fig 1B-D) (Garcez et al., 2017; Goto-Silva et al., 2021), which recapitulates the low expression levels of ACE2 mRNA observed in the human cortical tissue (Fig 1G). ACE2 levels are increased in patients with comorbidities, suggesting a relationship between ACE2 expression and susceptibility to severe COVID-19 (Pinto et al., 2020). Strategies to regulate the expression of ACE2 have been proposed as therapies to fight viral infection (Annweiler et al., 2020; Liu et al., 2020). Also, it remains unaddressed if neuropsychiatric conditions and neurodegenerative diseases impact the expression of ACE2 in the brain (Rocha et al., 2018) and how they could interfere with the susceptibility of CNS infection. Indeed, there is an increased risk for mental illness in COVID-19 patients, especially those who had previously psychiatric episodes (Mazza et al., 2020). Moreover, there is a raised concern that *in utero* infection might cause future neurodevelopmental disorders in the offspring (López-Díaz et al., 2021). In this sense, the current work contributes to the understanding of brain developmental consequences of SARS-CoV-2 by demonstrating that progenitors express relatively lower levels of ACE2 and CD147 and are not damaged by exposure to low MOIs, most likely to occur during a COVID pregnancy. In addition, human neurospheres and brain organoids could be a model for *in vitro* drug testing, including monitoring changes in ACE2 expression and correlation with susceptibility to SARS-CoV-2 infection.

One experimental aspect that drew our attention was the amount of virus used to infect cells in other studies. Previous reports have demonstrated SARS-CoV-2 antigens in NP and brain organoids (Mesci et al., 2020; Song et al., 2020; Zhang et al., 2020) using MOIs at least 100 times higher than the lowest MOI used in our investigation. Although productive infection in neural cells was claimed at MOI 10 (Zhang et al., 2020), we did not obtain infectious viral particles at similar experimental conditions (Fig 3A). Others observed viral proteins in a subset of neurons and SARS-CoV-2 sgRNA in the supernatant of these neuronal cultures, however, they did not gauged the production of virions (Dobrindt et al., 2021). This has been reviewed by Giani and Chen (2021) to show that SARS-CoV-2 is capable of entering neural cells (Giani and Chen, 2021). However, it remains possible that the viral variant and host cell genetics are determinant to permissiveness, i.e., presence of viral progeny, of SARS-CoV-2 infection in the brain.

The genetic comparison between our strain and the strains of other works showed little variations (Table 2) that, nevertheless, caused potentially significant alterations in the amino acid sequence of viral proteins (Table 3). Indeed, most of these variations might reflect the genetic diversity of the SARS-CoV-2 since the original outbreak (Phan, 2020). The strain used in the current work has a D614G mutation that renders the virus highly infectious by increasing ACE2 interaction (Korber et al., 2020). This D614G spike mutation has also been associated with increased binding of neutralizing antibodies (Weissman et al., 2020). Our strain has also a mutation in a cofactor of SARS-CoV-2 RNA-dependent RNA polymerase, nsp7, that did not lead to impairments in viral replication in permissive cells such as Vero (Fig 2C), however, this mutation could not be excluded as responsible for the differences observed between the studies. Finally, the T265I nsp2 mutation falls within the extracellular N-terminus of the protein (Angeletti et al., 2020) and is thought to cause structural changes, and which can now be found in all geographical locations (Rehman et al., 2020). Similarly to our work, Zhang and collaborators have also used a strain with increased sensitivity to neutralizing antibodies (Li et al., 2020). On the other hand, Song and collaborators used a viral strain that is not world spread (Alouane et al., 2020; Koyama et al., 2020; Laamarti et al., 2020) and it has been reduced in frequency over time since February 2020 (Alouane et al., 2020). In addition, the S5932F mutation on nsp14 (ExoN), could lead to altered proofreading exonuclease activity via weaker interaction with nsp10, facilitating the emergence of diverse viral sequences with potential selective advantage (Ogando et al., 2019).

Cells from lungs, intestine, heart and kidney, which have already been demonstrated to be sites of infection (Bradley et al., 2020; Mallapaty, 2020; Remmelink et al., 2020; Tabary et al., 2020), produce infectious viral progeny *in vitro* with MOIs ranging from 0.01 to 1 (Han et al., 2020; Lamers et al., 2020). Therefore, it seems unlikely that even in severe neuro-COVID cases such as the one cited here, neural cells would be exposed to quantities equal to or more than one plaque forming virus per cell. Our observation that ChP is infected to a much greater extent than other regions of the brain, together with the lack of infectious virus particles *in vitro* with MOI < 1, argue against the clinical relevance of using high titers of SARS-CoV-2 for modeling the infection *in vitro*. A study using viral titers of MOI 0.5 showed that SARS-CoV-2 exclusively infects ChP cells in brain organoids, but no other cell types. In the same report, even at higher viral titers (MOI 10), infection of neurons was minor (Pellegrini et al., 2020). Our observation that SARS-CoV-2 infects the ChP without significantly spreading to other brain areas (S1 Fig) and the absence of a productive infection in human NP with SARS-CoV-2 at MOI of 0.1, 1 and 10 even at increased inoculation time (Fig 3), corroborate these findings.

Despite non-permissiveness to SARS-CoV-2 infection of the brain parenchyma, the data presented here also points to direct pro-inflammatory and cytotoxic response of this tissue. This direct effect is probably related to the number of viral particles able to first get in contact with the CNS. It was observed in NP an increase in IL-7 and TNFα levels following infection with low MOIs, but not of other inflammatory cytokines such as IL-6. IL-7 has antiviral activity (Pellegrini et al., 2011) and neurotrophic effects in neurons (Araujo and Cotman, 1993; Michaelson et al., 1996, p. 7) and its increase at MOIs 0.01 and 0.1 may correlate with a protective neuroimmune response dependent on virus titer amount. TNF-α is secreted by astrocytes and microglia in response to injury, including viral infection, inducing neuronal damage (Figueiredo et al., 2019). It also regulates the permeability of the blood-brain barrier and neuronal plasticity (McCoy and Tansey, 2008; Probert, 2015). Both cytokines are increased in ICU-committed COVID-19 patients when compared to non-ICU patients, suggesting they are indeed correlated with disease severity (Huang et al., 2020). Certainly, these results are interesting with respect to a direct inflammatory response from neural cells that had been in contact with SARS-CoV-2 and deserve further investigation in neural tissue culture models that present well stablished immune response. A pro-inflammatory response in NP was suggested as a consequence of the upregulation of innate immune response protein TLR4 by ZIKA infection (Garcez et al., 2017). However, the immune response in NP still needs to be further comprehended in terms of how it parallels developing and mature human brains’ responses. Others have observed enrolment of pro-inflammatory pathways using different iPSC-derived models in contact with the new coronavirus (Makovoz et al., 2020; Giani & Chen, 2021). Nevertheless, the fact that proinflammatory cytokines were upregulated, however immature or unknown the NP immune response is, certainly merits mention and more studies. It is expected that in the complex scenario of the reaction to viral infection, multiple cytokines are regulated promoting cell protection and also limiting infection by inducing cell death. Therefore, the extent of neural damage should correspond to a balance between these responses.

We hypothesize that virus particles reaching the brain parenchyma could trigger inflammatory response leading to tissue damage. Notably, neurological symptoms are more frequent in severe patients with exacerbated inflammation, compared to mild or moderate ones, suggesting that an overall increase in the inflammatory response may set place for CNS damage (Mao et al., 2020). Low type I interferon response in infants’ brains may correlate with higher susceptibility for virus entry in CNS, but also patients with previous neurological and psychiatric disorders which may have weakened immune response or enhanced inflammation should be closely monitored (Mazza et al., 2020; Singh et al., 2020).

## 5. Acknowledgments

The authors would like to thank Claudia Figueiredo and Claudio Ferrari for helpful discussions during the elaboration of this manuscript; and Fernando Colonna Rosman and Leila Maria Cardão Chimelli for their support in the tissue’s analysis.

