## Supplementary Material for "Non-permissive SARS-CoV-2 infection in human neurospheres"

**Supplementary Figures and Tables**

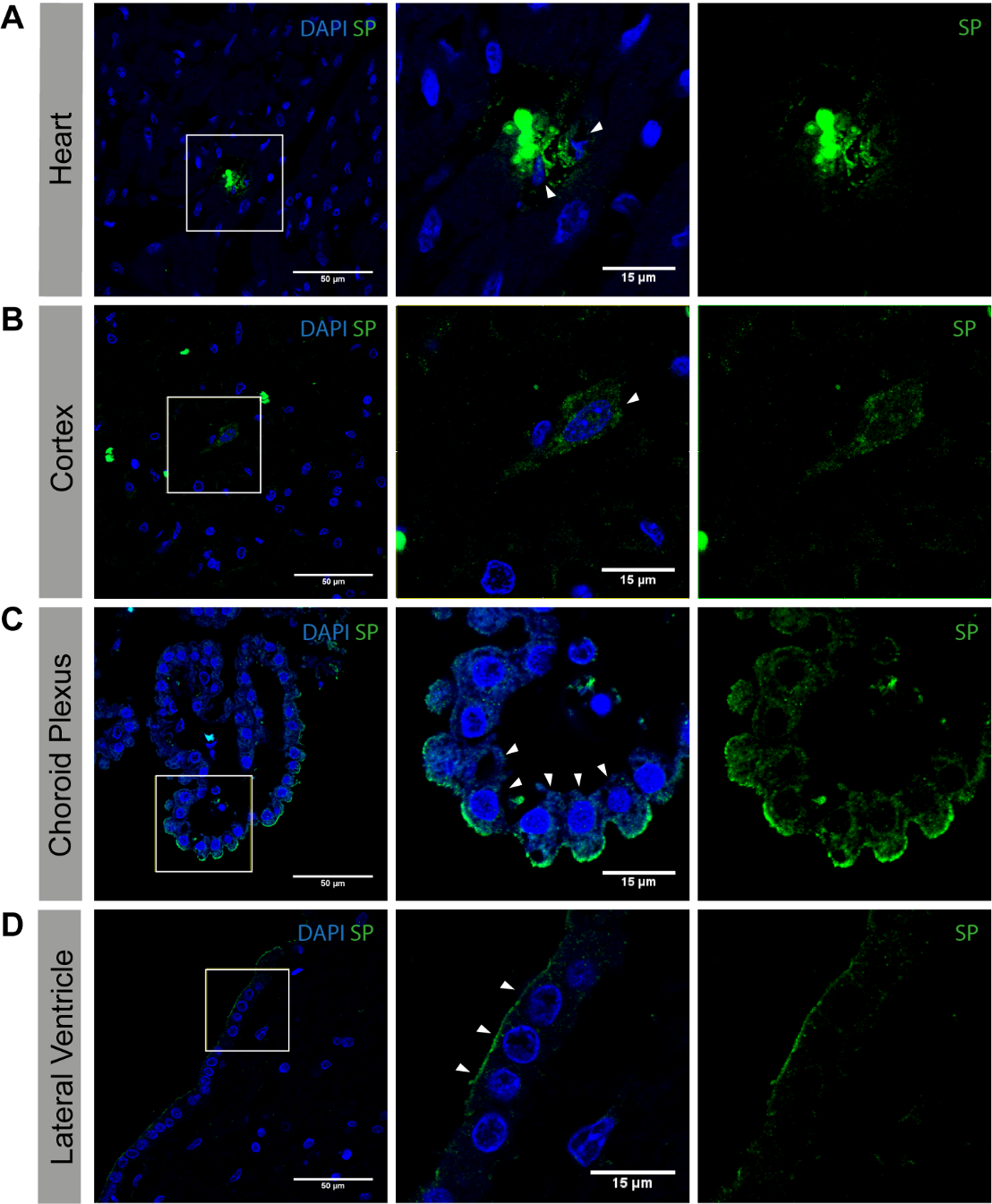

**S1 Fig**. **SARS-CoV-2 identified in the heart, cerebral cortex, choroid plexus (ChP) and lateral ventricle of a child perished from COVID-19.** IF staining in the heart (A), cortex (B), ChP (C) and LV (D) tissues with anti-SARS-CoV-2 spike protein (SP). Calibration bar: 50 µm for the upper panel and 15 µm for the lower panel. Complete pathological analysis was detailed in [1].

**S1Table. SARS-CoV-2 detection in human samples by RT-qPCR.**

| **Specimen Type** | **2019-nCoV_N1**  **Assay** | **2019-nCoV_N2**  **Assay** | **Hs_RPP30** | **Result Interpretation** |
| --- | --- | --- | --- | --- |
|  | **(Copies/Reaction)** | **(Copies/Reaction)** | **(Cp mean)** | **(Detected/Total Tested)** |
| **Lung** | 505 | 1,219 | 35.7 | 2019-nCoV  detected (3/3) |
|  | 6,201 | 2,019 | 36.4 |  |
|  | 524 | 679 | 31.5 |  |
| **Cortex** | 202 | 364 | 28.7 | 2019-nCoV  detected (2/2) |
|  | 1,162 | 5,631 | 26.9 |  |
| **Heart** | 100 | 652 | 25.1 | 2019-nCoV  detected (4/5) |
|  | 99 | 491 | 26.0 |  |
|  | ND | 4 | 30.3 |  |
|  | 2 | 11 | 34.3 |  |
|  | 2 | 4 | 35.8 |  |
| **Choroid Plexus** | ND | 29 | 32.4 | 2019-nCoV  detected (1/1) |
| **Lateral Ventricle** | ND | 9 | 34.1 | 2019-nCoV  detected (1/3) |
|  | ND | 25 | 34.4 |  |
|  | 3 | 32 | 36.4 |  |
| **2019-nCoV_N**  **(Positive Control)** | 880 | 1,720 | Undetermined | 2019-nCoV  detected (4/4) |
|  | 843 | 1,747 | Undetermined |  |
|  | 792 | 1,884 | Undetermined |  |
|  | 718 | 2,233 | Undetermined |  |
| **Hs_RPP30**  **(Positive Control)** | Undetermined | Undetermined | 28.2 | 2019-nCoV not detected (4/4) |
|  | Undetermined | Undetermined | 28.3 |  |
|  | Undetermined | Undetermined | 28.3 |  |
|  | Undetermined | Undetermined | 27.6 |  |
| **No template Control (NTC)** | Undetermined | Undetermined | Undetermined | 2019-nCoV not detected (4/4) |
|  | Undetermined | Undetermined | Undetermined |  |
|  | Undetermined | Undetermined | Undetermined |  |
|  | Undetermined | Undetermined | Undetermined |  |
| **Human Specimen Control**  **(Child lung biopsy –**  **Negative Control)** | Undetermined | Undetermined | 24.2 | 2019-nCoV not detected (2/2) |
|  | Undetermined | Undetermined | 23.2 |  |
| **Human Specimen Control (Human iPSC-derived Astrocytes - Negative Control)** | Undetermined | Undetermined | 19.4 | 2019-nCoV not detected (4/4) |
|  | Undetermined | Undetermined | 18.5 |  |
|  | Undetermined | Undetermined | 18.8 |  |
|  | Undetermined | Undetermined | 19.4 |  |
| **Human Specimen Control (Adult Nasopharyngeal Swab - Negative Control)** | Undetermined | Undetermined | 27.2 | 2019-nCoV not detected (4/4) |
|  | Undetermined | Undetermined | 27.3 |  |
|  | Undetermined | Undetermined | 31.9 |  |
|  | Undetermined | Undetermined | 31.6 |  |
| **Human Specimen Control (Adult Nasopharyngeal Swab - Positive Control)** | 68,183 | 93,336 | 26.6 | 2019-nCoV  detected (4/4) |
|  | 76,97 | 93,694 | 26.6 |  |
|  | 2,832,498 | 4,346,556 | 28.9 |  |
|  | 11,184,843 | 15,804,913 | 27.2 |  |

The absolute quantification (number of copies/reactions) of N1 and N2, and standard deviations of crossing points (Cp) values of RPP30 were calculated from data obtained in all analyzed distinct fragments from the same post-mortem tissue specimens of heart (n = 5), choroid plexus (n = 1), lateral ventricle (n = 3), and cortex (n = 2). Post-mortem lung sample derived from a non-Covid-19 child (n=2), iPSC-derived astrocytes (n=4) and non-Covid-19 adult nasopharyngeal swabs samples were used as negative control. 2019-nCoV_N, Positive Control plasmid; Hs_RPP30, Homo sapiens ribonuclease P/MRP subunit p30; NTC, no template control; ND, not determined.

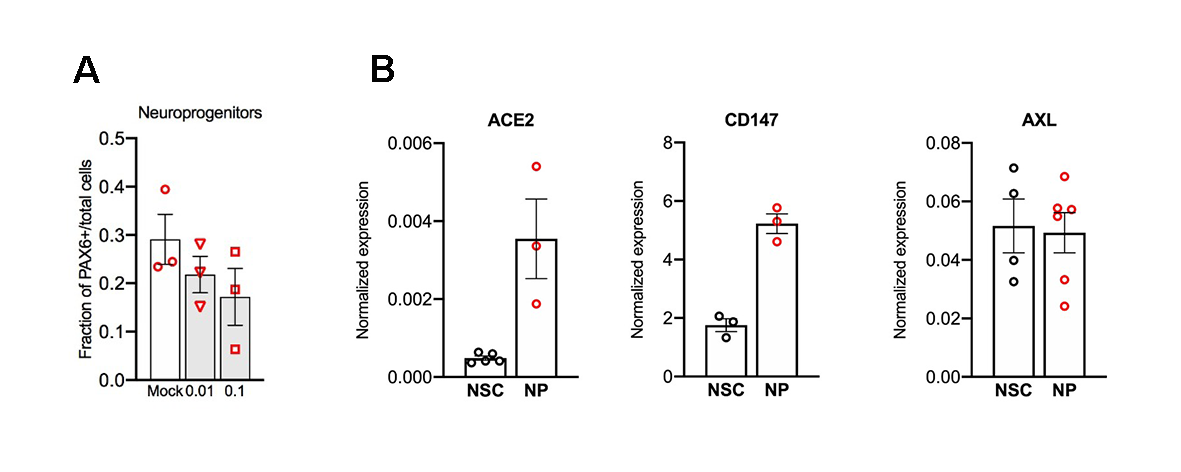
**S2 Fig. Analysis of neural progenitor population in SARS-CoV-2 infected neurospheres and expression of SARS-CoV-2 receptors in this model.** (A) Quantification of neural progenitor (PAX6+) population at MOI 0.01 and 0.1, 5 d.p.i., from 3 cell lines (Table 1) from one experimental infection. Quantification of progenitor nuclei from the NP was obtained by isotropic fractionation and immunostaining for PAX6, divided by total nuclei (DAPI+). (B) Comparison of SARS-CoV-2 receptors expression from neural stem cells (NSC) and neurospheres (NP) by qRT-PCR.

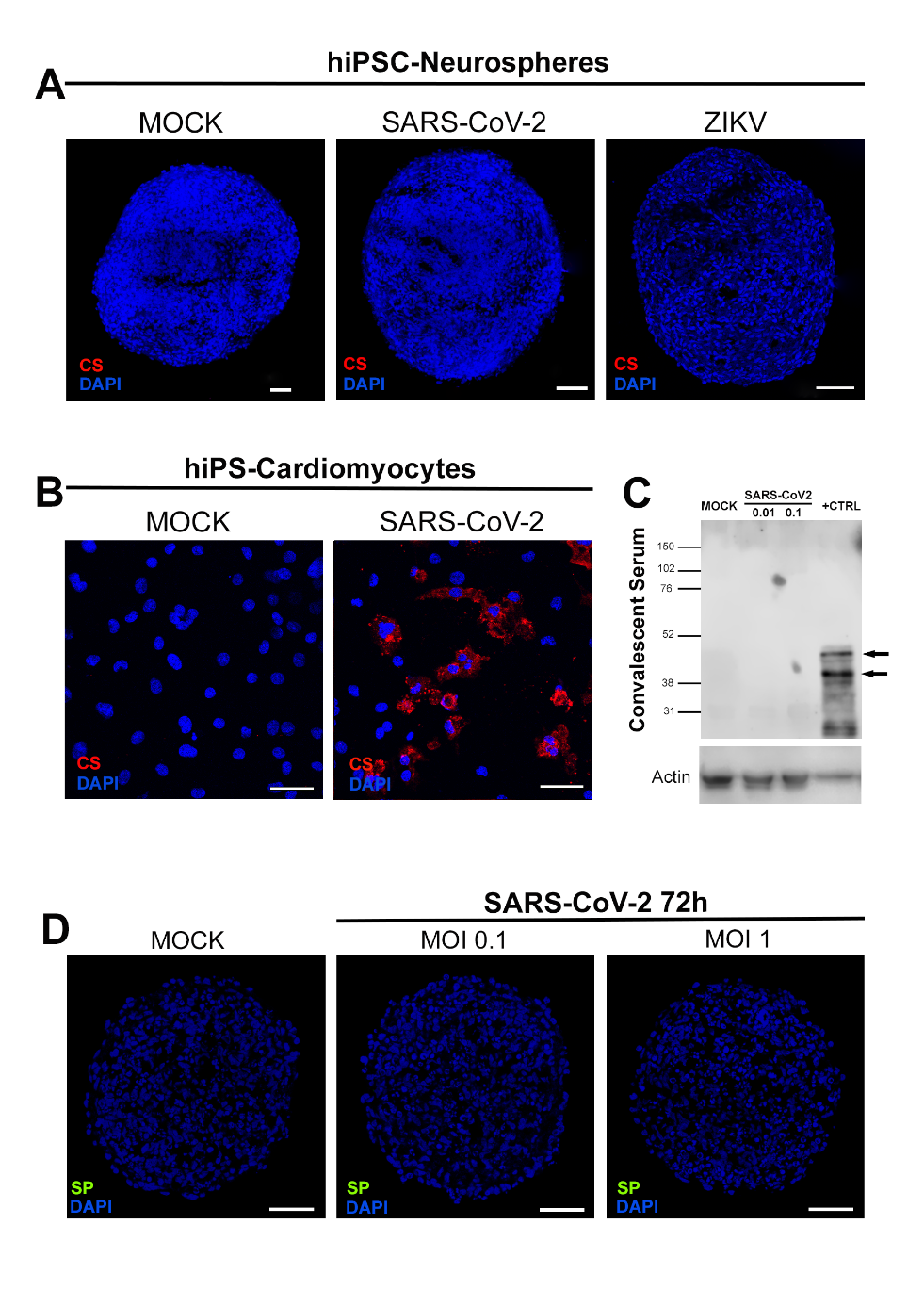

**S3 Fig. SARS-CoV-2 and Zika virus (ZIKV) infection in neurospheres and Vero cells**. Human neurospheres were infected with SARS-CoV-2 (MOI 0.1 for 1h, 5 d.p.i.) or ZIKV (MOI 0.5 for 2h, 3 d.p.i), followed by fixation in PFA 4% for microscopic analysis. IF staining for anti-SARS-CoV-2 CS (red) in (A) neurospheres cryosections and (B) SARS-CoV-2-infected cardiomyocytes (MOI 0.1 for 1h, 3 d.p.i). Nuclei were counterstained with DAPI (blue). Calibration bar: 50 µm (C) Detection of SARS-CoV-2 proteins by western blotting using anti-SARS-CoV-2 CS. Protein extracts from Vero cells (MOI 0.1 for 1h, 2 d.p.i) were used as positive controls. Gel loading was assessed with beta-actin staining. (D) Human neurospheres were exposed to SARS-CoV-2 (MOI 0.1 and 1 for 24h, 72 hours post infection) followed by fixation in PFA 4% for microscopic analysis. IF staining for anti-SARS-CoV-2 SP (green) in neurospheres cryosections. Nuclei were counterstained with DAPI (blue). Calibration bar: 40 µm.

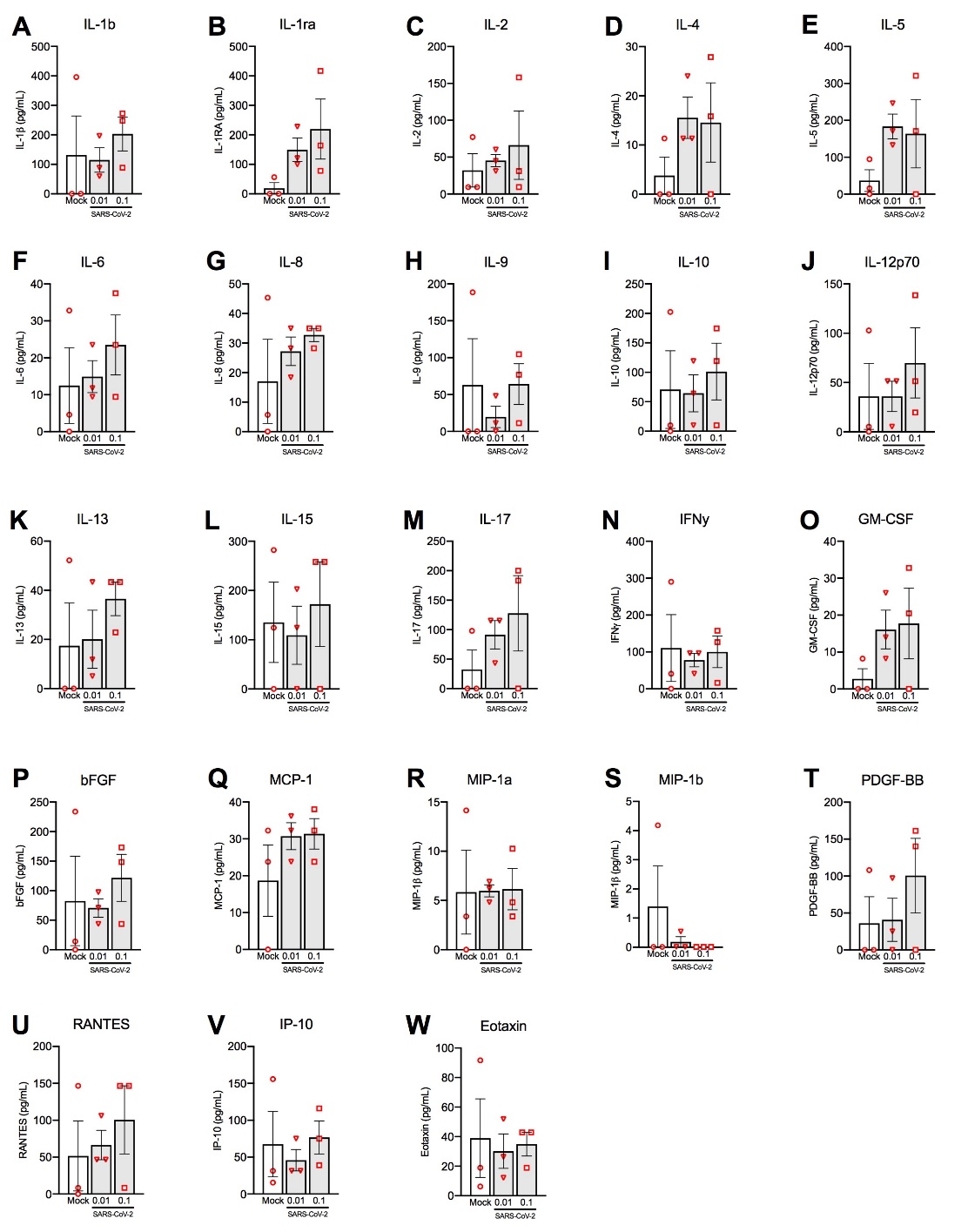

**S4 Fig. Multiplex Luminex analysis of cytokine and chemokine in SARS-CoV-2-infected neurospheres.** Human neurospheres were exposed to SARS-CoV-2 (MOI 0.01 or 0.1) for 1h, followed by analysis of the supernatant 5 days post-infection.

**Supplementary Material and Methods**

**Biological specimens and ethical statements**

Specimens from heart and brain (frontal cortex, putamen/globus pallidus, temporal lobe, lateral ventricle, choroid plexus (ChP), cortex, midbrain, pons and medulla oblongata) from a one-year-old child deceased due to COVID-19 were obtained after a signed informed consent by the legal guardians. The study was submitted to the internal Ethics Committee approval (CAAE number: 37211220.0.0000.5249) and fully conveyed as a case report by Gomes and collaborators [1]. The convalescent serum (CS) was obtained from recovered COVID-19-patients, after institutional review board approval (CAAE number: 30650420.4.1001.0008), as part of a national surveillance program.

**Sample preparation and total RNA isolation from human specimens**

Post-mortem tissue specimens from heart and brain (frontal cortex, lateral ventricle, and choroid plexus) were sliced into thick sections. Then, the samples were homogenized with 3.0 mm TriplePure Zirconium homogenizer beads (Benchmark Scientific, New Jersey, EUA) using the BeadBugTM Microtube Homogenizer apparatus (D1030-E, Benchmark Scientific). The total RNA was isolated with TRIzol™ Reagent (Thermo Fisher Scientific, Massachusetts, EUA), according to the manufacturer’s instructions.

For the negative controls, total RNA was isolated from 140 μL of respiratory specimens, obtained through nasopharyngeal swabs from upper respiratory tract, from four healthy adults using the QIAamp® Viral RNA Mini Kit (52906, Qiagen, Hilden, Germany), following manufacturer's instructions. Total RNA from Human iPSC-derived astrocytes were extracted using PureLinkTM RNA Mini Kit (12183018A, Thermo Fisher Scientific), according to manufacturer's protocol. RNA concentration and quality were quantified using a NanoDrop 2000 spectrophotometer (Thermo Fisher Scientific).

**Virus evaluation in human specimens**

RT–qPCR was performed on each sample using the 2019–nCoV CDC RUO Kit (IDT: 10006713) and 2019–nCoV CDC RUO Primers and Probes (PN: 10006713) for the detection of viral RNA (SARS-CoV-2 nucleocapsid N1 and N2 fragments) and the RNase P (RP) primer set for the detection of human RNase P RNA (Integrated DNA Technologies, Iowa, EUA). For each specimen, three separated reactions were set up in a 96-well plate including N1, N2, and RP primers and probes. RT-qPCR was carried out in with a total reaction volume of 20 µL containing 15 µL of GoTaq® Probe 1-Step RT-qPCR System (A6120, Promega, Wisconsin, EUA) comprised of the following components: 3.1 µL ultrapure water, 10 µL GoTaq® Probe qPCR Master Mix with dUTP (2X), 0.4 µL GoScriptTM RT Mix for 1-Step RT-qPCR, 1.5 µL primer/probe sets for either N1, N2, or RP (IDT) and 5 µL of extracted RNA.

To monitor assay performance, all reactions were carried out with negative controls (human specimen controls: human iPSC-derived astrocytes; a post-mortem lung tissue of a non-COVID four-month-old infant, who died due to respiratory failure caused by bilateral pneumonitis; adult nasopharyngeal swabs negatives for respiratory viruses; and no RNA template control with UltraPure DNase/RNase-Free Distilled water) and positive controls [2019-nCoV_N Positive Control plasmid, IDT: 10006625 and Hs_RPP30 (Homo sapiens ribonuclease P/MRP subunit p30) Positive Control, IDT: 10006626) and adult nasopharyngeal swabs positives for COVID], which were incorporated into each run to ensure proper testing control. Briefly, the reactions were performed on a StepOnePlusTM Real-Time PCR System thermocycler (Thermo Fisher Scientific). Thermal cycling conditions comprised a holding stage at 45 °C for 15 min, 95 °C for 2 min, followed by 45 cycles of denaturation at 95 °C for 15 seconds, and annealing and extension at 60 °C for 1 minute. The RT-qPCR results were analyzed according to “FDA Inform Diagnostics SARS-CoV-2 RT-PCR Assay”, available in https://www.fda.gov/media/139572/download.

The standard curve for virus quantification was prepared with 10-fold serial dilutions (ranging from 1 x 10^6^ to 10 copies/reaction) of synthetic viral RNA in a matrix containing pooled RNA isolated from four human respiratory samples from nasopharyngeal swabs negatives for respiratory viruses. Linear regression analysis was performed with the N1 and N2 assays from US CDC real-time reverse transcription PCR panel. Each dilution point was run in triplicates.

**Immunofluorescence staining**

Specimens from heart and brain (frontal cortex, putamen/globus pallidus, temporal lobe, lateral ventricle, choroid plexus (ChP), cortex, midbrain, pons and medulla oblongata) were fixed in 10% neutral buffered formalin. After 48 hours, part of the samples was dehydrated in a series of increasing concentrations of ethanol, cleared in xylol and paraffin embedded. For immunofluorescence staining, a tissue microarray (TMA) from the paraffin blocks of the above-mentioned tissues was produced by the protocol described by Pires and collaborators [2] with modifications. Some of the remaining fixed tissues were maintained in 70% ethanol before being further processed for total RNA isolation. 4 µm-sections of TMA were dewaxed with xylene and hydrated with decreasing concentrations of ethanol. Then, sections were incubated with 10 mM citrate buffer (pH 6.0) for 30 minutes for antigenic recovery, followed by incubation in permeabilization/blocking solution (0.3 % Triton X-100/ 3% bovine serum albumin, BSA, in PBS) for 2 hours. The sections were incubated with primary antibody anti-SARS-CoV-2 spike protein monoclonal antibody (SP) (1:500, G632604 - Genetex) at 4°C overnight. Next, sections were incubated with the secondary antibody goat anti-mouse AlexaFluor 488 (1:400, A-11008 - Invitrogen) for 1 hour. Nuclei were counterstained with 0.5 µg/mL 4′-6-diamino-2-phenylindole (DAPI) and the slides were mounted with Aqua-Poly/Mount (Polysciences).

Neurospheres were fixed in 4% paraformaldehyde solution (Sigma-Aldrich) for 1h, followed by cryopreservation with 30% sucrose solution overnight. Then, samples were embedded in O.C.T compound (Sakura Finetek, Netherland) and frozen at -80 ˚C. The O.C.T blocks were sectioned at 20 μm-slices with a Leica CM1860 cryostat. After washing with PBS, sections were incubated in permeabilization/blocking solution (0.3% Triton X-100/ 3% goat serum) for 2h. The primary antibodies were incubated overnight at 4˚C. For SARS-CoV-2 detection was used anti-SARS-CoV-2 convalescent sera (CS) (1:1000) or anti-SARS-CoV-2 SP (1:500, Cat#ab272420 - Abcam). For NP characterization were used: anti-Nestin (1:2000, Cat#RA22125 - Neuromics), anti-S100b (1:100, Cat#ab52642 - Abcam), anti-MAP-2 (1:250, Cat#M1406 - Sigma). Then, sections were incubated with secondary antibody goat anti-human Alexa Fluor 647 (A21445 - Thermo Fisher Scientific), goat anti-rabbit Alexa Fluor 488 (A-11008 – Invitrogen) or goat anti-mouse Alexa Fluor 488 (A-11001 – Invitrogen); all 1:400, for 2h. Nuclei were counterstained with DAPI for 10 minutes and the slides were mounted with Aqua-Poly/Mount.

After fixation, Cardiomyocytes were washed with PBS and then incubated with permeabilization/blocking solution (0.3% Triton X-100 / 3% bovine serum albumin) for 1h. Then, cells were incubated with primary antibody overnight at 4º. For SARS-CoV-2 detection was used anti-SARS-CoV-2 CS (1:1000). For CM characterization was used anti-cardiac troponin T (TNNT) (1:2500, Cat#MA5-12960 – Invitrogen). Day after, cells were incubated with the secondary antibody goat anti-Human Alexa Fluor 647 (A-21445 - Invitrogen) or goat anti-mouse Alexa Fluor 488 (A-11001 – Invitrogen); both 1:400; for 1h. Nuclei were counterstained with DAPI for 5 minutes and mounted with 50% PBS-Glycerol.

Images of neurospheres and cardiomyocytes were acquired on a Leica TCS-SP8 confocal microscope with the 63x and 20x objective, respectively.

**Identification of neural progenitor population**

Nuclei from human neurospheres were obtained by isotropic fractionation (Herculano-Houzel and Lent, 2005). Samples containing nuclei were submitted to antigen retrieval with citrate buffer (10 mM Sodium citrate, 0.05% Tween 20, pH 6.0) at 92 ^o^C for 20 min and plated in 96-well plates coated with 0.1 mg/ml poly-L-lysine. Nuclei were immunostained with anti-PAX6 antibody as follows: first they were permeabilized with 0.3% Triton X-100 in PBS for 20 min, then washed 3 times with PBS, blocked with BSA 3% in PBS for 1 h and incubated with anti-PAX6 (1:200, KloneLife sc-11357 rabbit) antibody diluted in BSA 3% in PBS overnight. Subsequently, nuclei were washed 3 times with PBS and incubated with secondary antibody goat anti-rabbit Alexa Fluor 488 (1:400) for 1h. Then the nuclei were washed 3 times with PBS and counterstained with DAPI for 5 minutes and 50% PBS-Glycerol was added to the wells. Image acquisition was carried out in an Operetta high-content imaging system with a 40x objective and high numerical apertures (NA) (PerkinElmer, USA). The data were analyzed using the high-content image analysis software Harmony 5.1 (PerkinElmer, USA). Five independent fields were evaluated from duplicate wells per experimental condition. Statistics analysis was made using one way ANOVA (significance set at p<0.05).

**Total RNA isolation and qRT-PCR**

Total RNA isolation of NP was performed using ReliaPrep^TM^ RNA Tissue Miniprep System (Promega Corporation) according to manufacturer’s instructions. For NSCs, total RNA was isolated using PureLink^TM^ RNA Mini Kit (Thermo Fisher Scientific). RNA concentration and quality were quantified on a NanoDrop^TM^ 2000c Spectrophotometer (Thermo Fisher Scientific); and integrity and purity were evaluated by 1.8% agarose gel electrophoresis using a photo documentation device equipped with a UV lamp (L-PIX, Loccus Biotecnologia). Then, samples were digested with DNase I, Amplification Grade, following the manufacturer’s instructions (Invitrogen, Thermo Fisher Scientific). 2 μg of RNA from DNAse-treated samples were reverse transcribed using M-MLV for complementary DNA generation (cDNA) (Thermo Fisher Scientific).

qRT-PCR for gene expression analysis of CD147 and AXL was performed as described in the Materials and Methods section of the manuscript, using the corresponding primers (CD147, forward: 5’-ACGTCCTGGATGATGACGAC-3’, reverse: 5’-GAAGAGTTCCTCTGGCGGAC-3’; AXL, forward: 5’-CCGTGGACCTACTCTGGCT-3’, reverse: 5’-CCTTGGCGTTATGGGCTTC-3’)
